# The energetic and allosteric landscape for KRAS inhibition

**DOI:** 10.1101/2022.12.06.519122

**Authors:** Chenchun Weng, Andre J. Faure, Ben Lehner

## Abstract

Thousands of proteins have now been genetically-validated as therapeutic targets in hundreds of human diseases. However, very few have actually been successfully targeted and many are considered ‘undruggable’. This is particularly true for proteins that function via protein-protein interactions: direct inhibition of binding interfaces is difficult, requiring the identification of allosteric sites. However, most proteins have no known allosteric sites and a comprehensive allosteric map does not exist for any protein. Here we address this shortcoming by charting multiple global atlases of inhibitory allosteric communication in KRAS, a protein mutated in 1 in 10 human cancers. We quantified the impact of >26,000 mutations on the folding of KRAS and its binding to six interaction partners. Genetic interactions in double mutants allowed us to perform biophysical measurements at scale, inferring >22,000 causal free energy changes, a similar number of measurements as the total made for proteins to date. These energy landscapes quantify how mutations tune the binding specificity of a signalling protein and map the inhibitory allosteric sites for an important therapeutic target. Allosteric propagation is particularly effective across the central beta sheet of KRAS and multiple surface pockets are genetically-validated as allosterically active, including a distal pocket in the C-terminal lobe of the protein. Allosteric mutations typically inhibit binding to all tested effectors but they can also change the binding specificity, revealing the regulatory, evolutionary and therapeutic potential to tune pathway activation. Using the approach described here it should be possible to rapidly and comprehensively identify allosteric target sites in many important proteins.

## Introduction

The GTPase KRas (KRAS) is somatically mutated in ∼10% of all cancers, including ∼90% of pancreatic adenocarcinoma, ∼40% of colorectal adenocarcinoma, ∼35% of lung adenocarcinoma and ∼20% of multiple myeloma ^1^. KRAS functions as an archetypal molecular switch, cycling between inactive GDP-bound and active GTP-bound states. The altered conformation and activity of KRAS upon GTP binding is an example of allostery, the long-range transmission of information from one site to another in a protein^2^. Many structures of KRAS have been determined, revealing major (but variable) rearrangements in the switch I and switch II regions that allow binding to effector proteins in GTP-bound states ^3^. KRAS effectors include the RAF proto-oncogene serine/threonine-protein kinase (RAF1 also known as CRAF) and Phosphatidylinositol 4,5-bisphosphate 3-kinase catalytic subunit gamma isoform (PIK3CG) and the signalling protein Ral guanine nucleotide dissociation stimulator (RALGDS). Guanine nucleotide exchange factors (GEFs) such as Son of sevenless homolog 1 (SOS1) catalyse release of GDP to activate KRAS whereas GTPase-activating proteins (GAPs) catalyse GTP hydrolysis to complete the cycle back to the inactive states. Cancer driver mutations interfere with this cycle, increasing the abundance of active GTP-bound effector-binding states ^4,5^.

Despite its identification as an oncoprotein 40 years ago ^6^, tens of thousands of scientific publications, and more than three hundred published structures of KRAS ^3^, only a single inhibitor of KRAS has been approved for clinical use: sotorasib, a covalent binder of one driver mutation, KRAS(G12C) ^7–9^. Sotorasib is an allosteric inhibitor that binds outside of the nucleotide and effector binding sites to lock KRAS(G12C) in inactive GDP-bound states, reducing effector binding and clinically validating the efficacy of allosteric KRAS inhibition ^7,9^. As for many medically important proteins, the development of therapeutics against KRAS is limited by the lack of information about inhibitory allosteric sites to target. Indeed, a comprehensive map of allosteric sites has not been generated for any oncoprotein or, indeed, for any disease target or any complete protein in any species.

Atlases of allosteric sites have the potential to greatly accelerate drug development, especially for the many human proteins considered ‘undruggable’ because of the lack of an appropriate active site or because they function via difficult to inhibit protein-protein interaction interfaces. In addition, among other benefits, allosteric drugs often have higher specificity than orthosteric drugs targeting conserved active sites ^10,11^.

### KRAS biophysics at scale

To comprehensively map inhibitory allosteric communication in KRAS, we applied our multidimensional deep mutational scanning approach ^12^. We used two rounds of nicking mutagenesis ^13^ to construct three libraries of KRAS variants in which every possible single amino acid (aa) substitution is present not only in the wild-type (WT) KRAS (4B isoform, aa1-188) but also in KRAS variants with a range of reduced activities (median of seven genetic backgrounds for each single mutant, Fig. 1a-d). Quantifying the effects of the same mutations in different genetic backgrounds provides sufficient data to infer the causal biophysical effects of each mutation (see below). In total, the library consists of >26,500 variants of KRAS, including >3,200 single aa substitutions and >23,300 double aa substitutions.

**Fig. 1.**
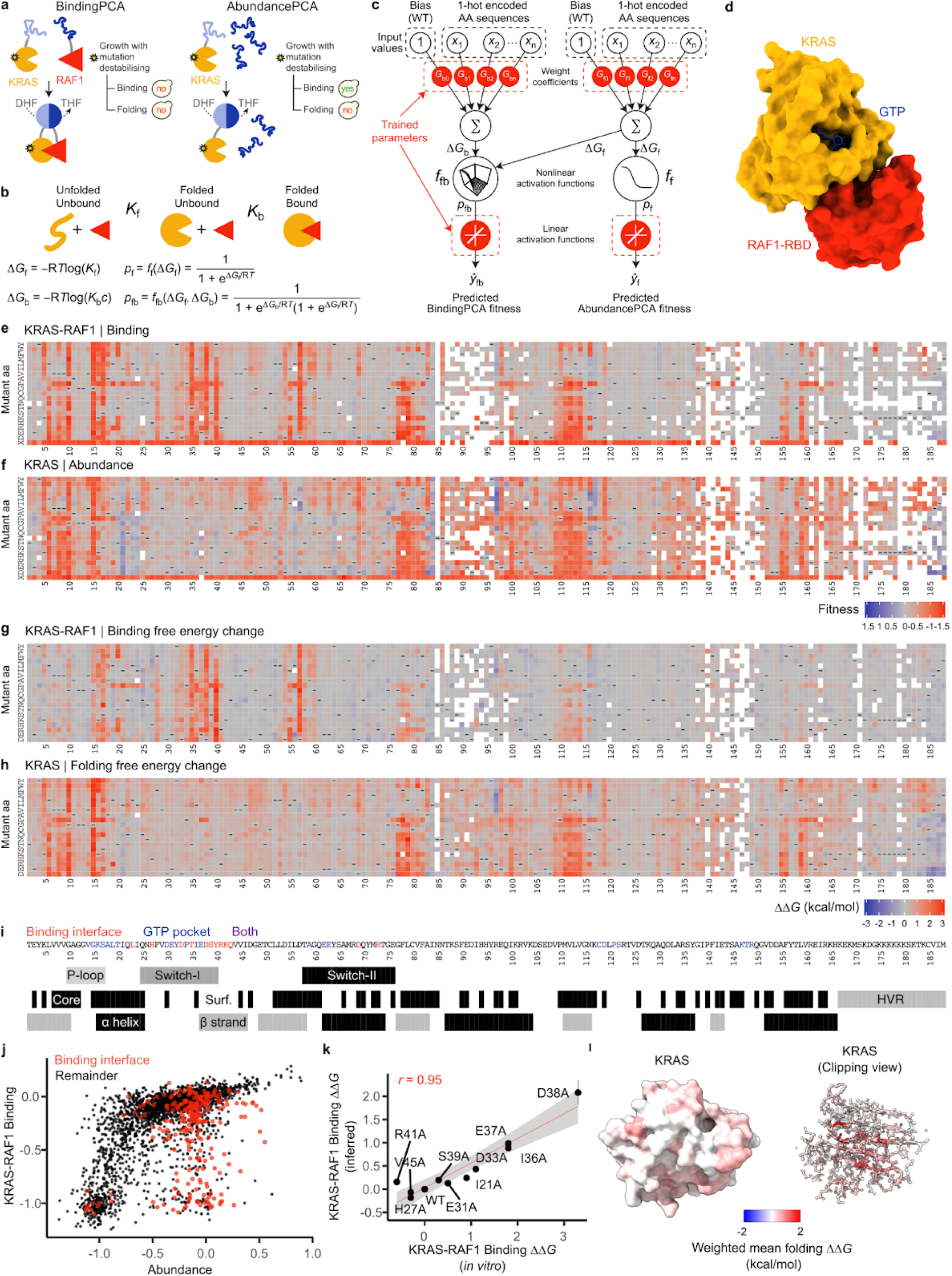
Mapping the energetic landscape of KRAS folding and binding to RAF1. **a**, Overview of ddPCA selections. yes, yeast growth; no, yeast growth defect; DHF, dihydrofolate; THF, tetrahydrofolate. **b**, Three-state equilibrium and corresponding thermodynamic model. Δ*G*_*f*_, Gibbs free energy of folding; Δ*G*_*b*_, Gibbs free energy of binding; *K*_*f*_, folding equilibrium constant; *K*_*b*_, binding equilibrium constant; *c*, binding partner concentration; *p*_*f*_, fraction folded; *p*_*fb*_, fraction folded and bound; *f*_*f*_, nonlinear function of Δ*G*_*f*_; *f*_*fb*_, nonlinear function of Δ*G*_*f*_ and Δ*G*_*b*_; R, gas constant; *T*, temperature in Kelvin. **c**, Neural network architecture used to fit thermodynamic models to the ddPCA data (bottom, target and output data), thereby inferring the causal changes in free energy of folding and binding associated with single amino acid substitutions (top, input values). **d**, 3D structure of KRAS bound to RAF1-RBD (Protein Data Bank (PDB) ID: 6VJJ). **e, f**, Heat maps of fitness effects of single aa substitutions for KRAS-RAF1 from BindingPCA (**e**) and AbundancePCA (**f**) assays. White, missing values; -, wild-type aa; X, STOP codon. **g, h**, Heat maps showing inferred changes in free energies of binding (**g**) and folding (**h**). **i**, Sequence and annotation of KRAS. Binding interface, RAF1 distance < 5 Å; GTP pocket, GTP or Mg2+ distance < 5 Å; core, relative accessible surface area (RSASA) < 0.25 (PDB:6VJJ). **j**, Scatter plot comparing abundance and binding fitness of single aa substitutions. Substitutions in the binding interface are indicated in red. **k**, Comparisons of model-inferred free energy changes to *in vitro* measurements^37^. Error bars indicate 95% confidence intervals from a Monte Carlo simulation approach (n = 10 experiments). Pearson’s *r* is shown. **l**, 3D structure (left) and clipping view (right) of KRAS with residues coloured by the weighted mean folding free energy change.

We first quantified the binding of these KRAS variants to the RAS-binding domain (RBD) of the oncoprotein effector RAF1. Binding was quantified using a protein-fragment complementation assay (BindingPCA) ^12,14,15^. Binding fitness was highly correlated among three independent replicate selections (Pearson’s *r* > 0.9, Extended Data Fig. 1a) and to previous data that used a different binding assay in a different cellular context (Pearson’s *r* = 0.82, Extended Data Fig. 1c ^16,17^.

As expected, mutations in the RAF1-binding interface strongly inhibit binding, as do variants in the nucleotide binding pocket (Fig. 1e,i). However, in total, 2,019 out of 3,231 single aa substitutions reduce binding to RAF1 (FDR = 0.05, two-sided *z*-test), including many outside of the interface including in the hydrophobic core of the protein (Extended Data Fig. 1d). This strongly suggests that many changes in binding to RAF1 are caused by changes in the abundance of folded KRAS, not by altered binding affinity ^12,18^.

### From molecular phenotypes to free energy changes

To disentangle the effects of mutations on KRAS folding and binding, we used a second selection assay, AbundancePCA ^12,19^, to quantify the cellular abundance of the KRAS variants. We refer to this combined approach of BindingPCA and AbundancePCA as ‘doubledeepPCA’ (ddPCA) ^12^. Plotting the RAF1 binding of each variant against its cellular abundance shows that many changes in binding can indeed be explained by reduced KRAS abundance (Fig. 1j). However, inspection of Figure 1j also reveals that a substantial number of variants have effects on binding that are much larger than can be accounted for by their reduced abundance, including many variants in the binding interface (red points in Fig. 1j).

Protein folding and binding relate to changes in the free energies of folding (Δ*G*_*f*_) and binding (Δ*G*_*b*_) by nonlinear functions derived from the Boltzmann distribution (Fig. 1b) ^12,18^. When mutational effects are combined additively in free energy they therefore cause non-additive changes in the molecular phenotypes of protein abundance and the abundance of a protein complex. The KRAS variant libraries contain a median of seven double mutants for each single aa substitution, providing sufficient experimental data to infer the causal free energy changes using MoCHI, a flexible package to fit mechanistic models to deep mutational scanning data using neural networks (see Methods) (Fig. 1c, Extended Data Fig. 1e). The fitted model predicts the mutant data extremely well (abundance median Pearson’s *r* = 0.86, binding median *r* = 0.96, Extended Data Fig. 1e) and the free energy changes are highly correlated with *in vitro* measurements (R =0.95, Fig. 1k).

### The RAF1 binding interface

In total, 2,241 out of 3,453 single aa substitutions are detrimental to folding and 843 out of 3,301 are detrimental to binding (FDR = 0.05, Fig. 1g, h). Mutations detrimental to folding are enriched in the hydrophobic core of the protein (odds ratio, OR = 1.92, Fisher’s exact test, *P* < 10e-16, Fig. 1h,l, Supplementary Movie 1). In contrast, mutations that increase the binding free energy are strongly enriched in the binding interface (OR = 6.02, *P* < 10e-16, Fig. 1g, 2a), with the mean absolute binding free energy changes upon mutation at each site identifying the binding interface (Fig. 2b,c, Supplementary Movie 2, receiver operating curve-area under curve, ROC-AUC = 0.8 compared to ROC-AUC = 0.65 when using the mean absolute binding fitness).

**Fig. 2.**
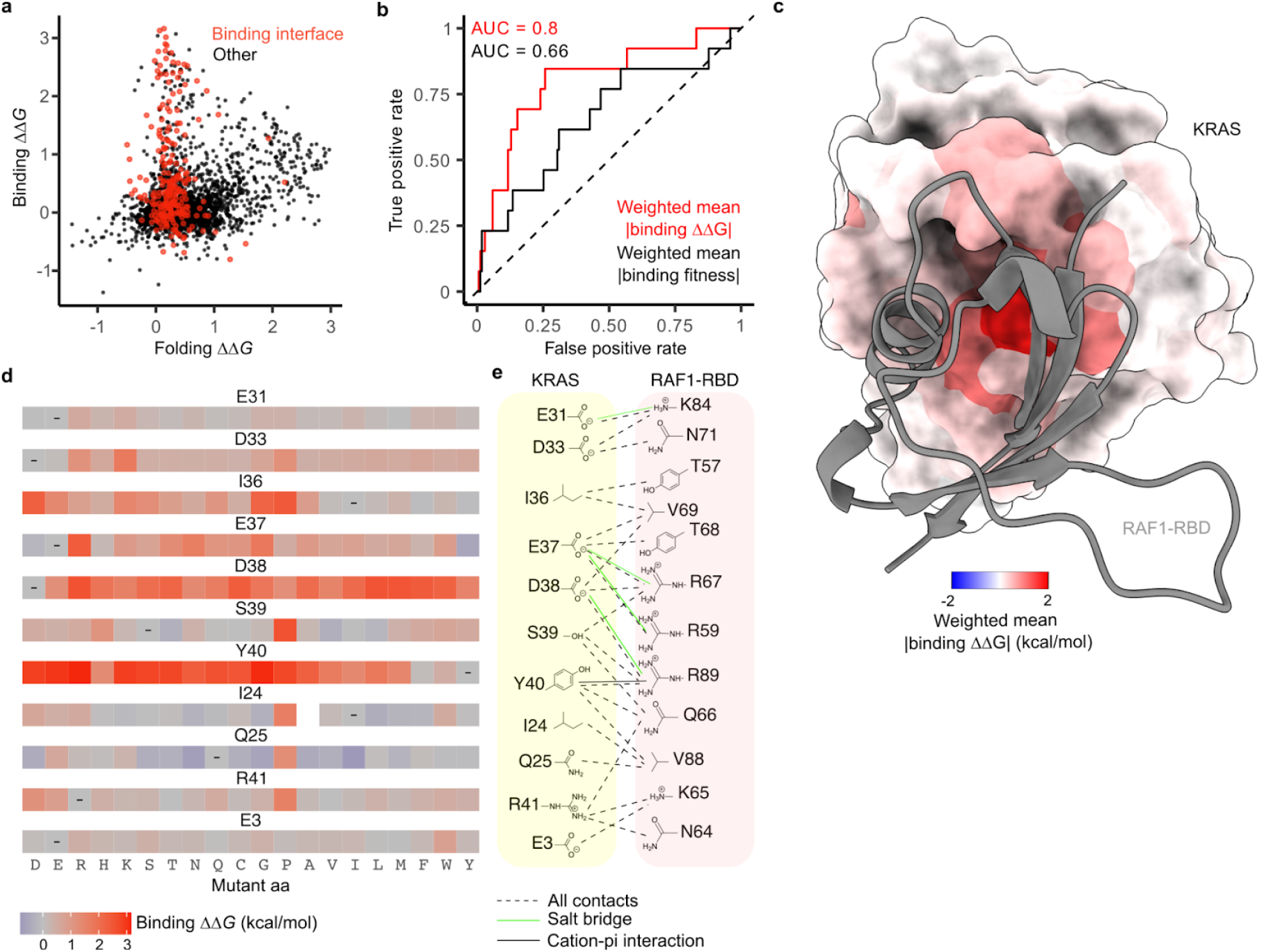
Free energy changes of mutations in the KRAS-RAF1 binding interface. **a**, Scatter plot comparing binding and folding free energy changes of single aa substitutions. **b**, ROC curves for predicting binding interface residues (RAF1 distance < 5 Å) using weighted mean absolute binding ΔΔ*G* (red) or using weighted mean absolute binding fitness (black). AUC = Area Under the Curve. **c**, 3D structure of KRAS bound to RAF1 in which residue atoms are coloured by the position-wise weighted mean absolute change in the free energy of binding to RAF1. RAF1-RBD is shown in grey. **d**, Heat maps of binding free energy changes in RAF1 binding interface residues. **e**, Direct contacts between KRAS and RAF1.

The interface residues most important for RAF1 binding include a mixture of charged (E37, D38) and hydrophobic (I36, Y40) residues. D38 cannot be changed to any other aa without detrimental effects on binding affinity, revealing a requirement for both negative charge and a particular side chain length at this site (Fig. 2d, e). In contrast, E37 can be replaced by D (shortening the side chain but retaining the negative charge) and also by Y, F or H, suggesting the salt bridge to RAF1 R67 can be replaced by an alternative interaction involving an aromatic side chain. Y40 can only be replaced by F, revealing the importance of the aromatic side chain which makes a cation-π interaction with RAF1 R89. I36 makes two hydrophobic contacts with RAF1, and whereas polar mutations at this position are detrimental, multiple hydrophobic substitutions are tolerated. Mutations at other residues that contact RAF1 are much better tolerated, indicating that these sites are less important for binding. For example, mutations at D33 tend to be mildly detrimental, with only charge-reversing mutations to R and K and mutation to P strongly inhibiting binding. Likewise, charge-reversing mutations and mutation to P are also most detrimental at R41, whereas mutations at the other two charged sites (E31 and E3) at the edge of the interface generally have little effect on the binding free energy.

### Inhibitory allosteric landscape for RAF1 binding

We next considered mutations outside of the binding interface. In total there are 361 distal mutations in 74 residues that increase the binding free energy to a greater extent than the average effect of mutations in the RAF1 binding interface (ΔΔ*G*_*b*_ greater than the weighted mean absolute binding free energy change of substitutions in binding interface residues, FDR = 0.05, Fig. 3a). Allosteric mutations defined in this manner are highly enriched in the physiological allosteric site of KRAS, the nucleotide binding pocket (157 mutations in 13 residues, OR = 7.68, Fisher’s exact test, *P* < 10e-16).

**Fig. 3.**
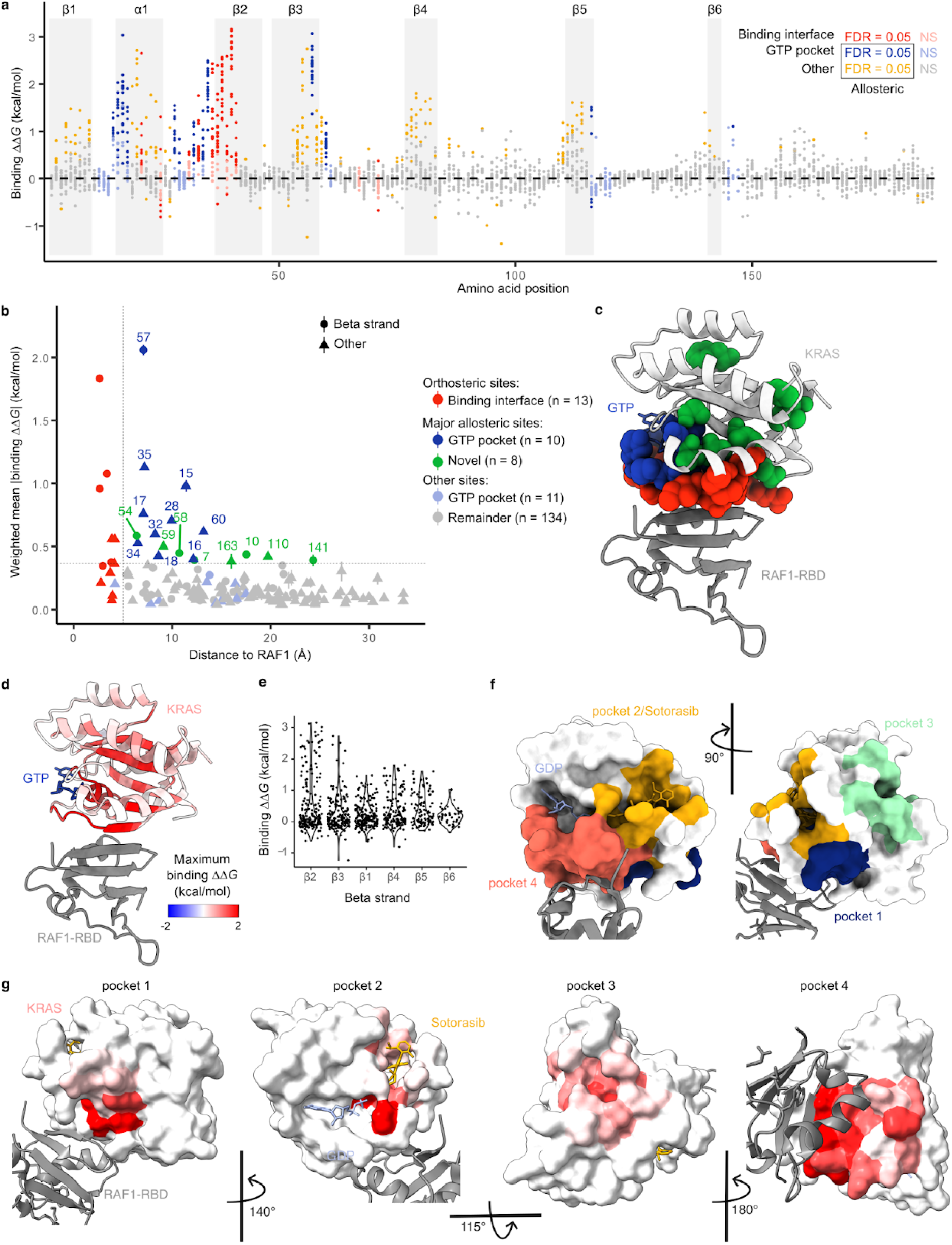
Allosteric regulation of KRAS binding to RAF1. **a**, Manhattan plot showing the binding free energy changes of all single aa substitutions. Points are coloured according to residue position and whether the corresponding binding ΔΔ*G* is significantly greater than the weighted mean absolute binding ΔΔ*G* of all mutations in the RAF1 binding interface (two-sided *z*-test, FDR = 0.05). **b**, Relationship between the position-wise average absolute change in free energy of binding to RAF1 and the minimal side chain heavy atom distance to RAF1. Major allosteric sites are defined as non-binding-interface residues with weighted mean absolute change in free energy of binding higher than the average of binding-interface residue mutations (horizontal dashed line). Error bars indicate 95% confidence intervals (n ≥ 10). **c**, 3D structure (PDB:6VJJ) of KRAS bound to RAF1 (grey) with binding interface and major allosteric site residue atoms of KRAS coloured as in **b. d**, Similar to c except KRAS residues are coloured by maximum binding ΔΔ*G*. **e**, Violin plot showing the decay of binding free energy change across successive strands in the beta sheet. Beta strands are ordered by increasing distance to RAF1 in the 3D structure. **f**, 3D structure (PDB:6OIM) of KRAS bound to GDP (blue), sotorasib (yellow) and RAF1 (grey) with KRAS surface coloured according to previously described pockets in KRAS (pocket 2, Sotorasib distance < 5 Å; pocket 1, 3, 4 ^20^). **g**, Similar to **h** except KRAS pockets are coloured by maximum binding ΔΔ*G*.

### Enhanced allosteric communication across a beta sheet

We first focussed on the residues where many different mutations have strong allosteric effects. Defining major allosteric sites as residues where the mean absolute change in binding free energy upon mutation is equal to or greater than that in binding interface residues identifies a total of 18 sites (Fig. 3b, c). 10 of these major allosteric sites are located in the physiological allosteric site - the nucleotide binding pocket (Fig. 3b, c). The additional 8 major allosteric sites are residues V7, G10, D54, T58, A59, P110, F141 and I163 (Fig. 3b). Three of these residues are close to the binding interface, with Asp 54 adjacent to the binding interface and Thr 58 and Ala 59 connecting the binding interface to the nucleotide binding pocket (Fig. 3c, Supplementary Movie 3).

Strikingly, 5 of the 8 novel major allosteric residues are located in the central beta sheet of KRAS (Fig. 3b, c). Within the beta sheet, the binding free energy changes are largest for mutations in residues in the first strand that contacts RAF1 and they progressively decrease in each subsequent strand of the sheet (Fig. 3d, Extended Data Fig. 2a-d, Supplementary Movie 4). This decay of the strength of allosteric effects across the sheet is consistent with local energetic propagations underlying allosteric communication. A similar, yet weaker, distance-dependent decay is observed for residues outside of the beta-sheet (Extended Data Fig. 3c). Propagation appears more efficient across the sheet than along the backbone within a strand, with residues in the first strand that do not contact RAF1 being depleted for allosteric mutations (Fig. 3a, Extended Data Fig. 2b, OR = 0.16, Fisher’s exact test, *P* = 1e-3). Allosteric communication therefore seems particularly effective across the central beta sheet of KRAS.

### KRAS has four allosterically active surface pockets

We next considered the effects of mutations in the surface residues of KRAS, focussing on the four previously described pockets in addition to the nucleotide-binding pocket (Fig. 3e, Supplementary Movie 5) ^20^.

Pocket 1 (also called the switch I/II pocket) is located behind switch II between the central beta sheet and alpha helix 2 and is the binding site for multiple inhibitors that are effective in pre-clinical models ^21,22^. Many mutations in pocket 1 allosterically inhibit RAF1 binding (57 mutations in 10 residues, FDR = 0.05, Fig. 3f, Extended Data Fig. 2e), consistent with the demonstrated ability of pocket 1 engagement to inhibit effector binding.

Pocket 2 (also called the switch II pocket) is located between switch II and alpha helix 3 and is the binding site of sotorasib, the clinically approved allosteric inhibitor of KRAS(G12C) ^23^. 71 mutations in 9 residues that contact sotorasib allosterically inhibit RAF1 binding (Figure 3g, Extended Data Fig. 2f). Thus, both mutations and small molecules binding to pocket 1 and pocket 2 can allosterically inhibit KRAS activity.

Pocket 3 of KRAS is located in the C-terminal lobe of the protein and is the most distant pocket from the RAF1 binding interface (Fig. 3e). The effects of pocket 3 engagement are not well described ^20^ and pocket 3 has received little attention for therapeutic development ^21^. However, our data reveal that pocket 3 is allosterically active, with 20 mutations in 6 residues in pocket 3 inhibiting binding to RAF1 (Fig. 3h, Extended Data Fig. 2g). Indeed helix 1 in pocket 3 is the only secondary structure element outside of the beta sheet enriched for allosteric mutations (Extended Data Fig. 2b, odd ratio = 5.7, Fisher’s exact test, *P* < 10e-16). Despite its distance from the effector binding interface, pocket 3 should be prioritised as a site for the development of KRAS inhibitors.

Finally, pocket 4, which is located immediately behind the flexible effector binding loop, contains 105 allosteric mutations in 9 residues that do not contact RAF1 (Fig. 3f, Extended Data Fig. 2h). Our data therefore validate all four surface pockets of KRAS as allosterically active, with perturbations in all pockets having large inhibitory effects on RAF1 binding. This strongly argues for the development of molecules targeting all four pockets as potential KRAS inhibitors.

### Energetic landscapes for six KRAS interactions

Like most oncoproteins, KRAS binds many different proteins as part of its physiological and disease-relevant functions ^2^. Many of these interaction partners bind a common surface of KRAS – the effector-binding interface – making KRAS an interesting model of multi-specificity in molecular recognition ^2^. To our knowledge, the effects of mutations on binding energies for multiple interaction partners have not been comprehensively profiled for any protein. Moreover, quantifying KRAS binding to multiple interaction partners provides an opportunity to quantify the conservation and specificity of allosteric effects in a signalling hub (Fig. 4a).

**Fig. 4.**
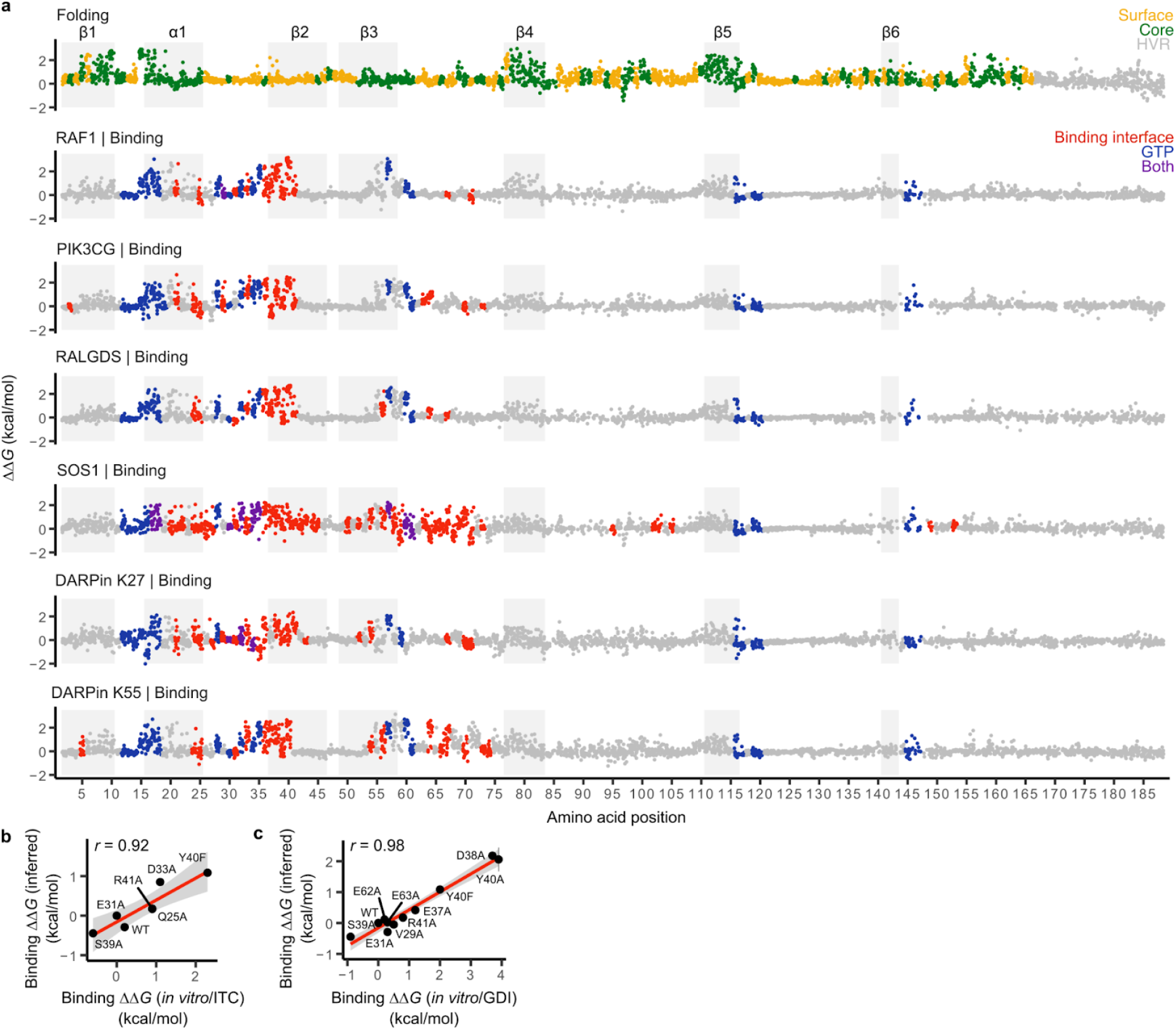
Seven KRAS free energy landscapes. **a**, Manhattan plots showing the folding and binding free energy changes of all single aa substitutions. Dark grey rectangles indicate beta strands, light grey rectangles indicate alpha helix 1; HVR, hypervariable region. Binding interface, indicated binding partner distance < 5 Å (RAF1, PDB:6VJJ; PIK3CG, PDB:1HE8; RALGDS, PDB:1LFD; SOS1, PDB:1NVW; DARPin K27, PDB:5O2S; DARPin K55, PDB:5O2T). **b, c**, Comparisons of binding free energy changes to *in vitro* measurements. Pearson’s *r* is shown. Error bars indicate 95% confidence intervals from a Monte Carlo simulation approach (n = 10 experiments).

We quantified the binding of the >26,000 KRAS variants to six interaction partners: the three KRAS effector proteins RAF1, PIK3CG and RALGDS, the GEF SOS1 and two DARPins, K27 and K55, synthetic antibody-like molecules selected to bind GDP- and GTP-bound KRAS, respectively. The structures of all six complexes have been determined ^24–28^.

The data for all six binding selections were highly reproducible (Extended Data Fig. 1a, 3a), and we used MoCHI to simultaneously fit a thermodynamic model to the molecular phenotypes of the variants in all seven experimental datasets (see Methods, Extended Data Fig. 3b). Each single aa change in KRAS therefore has seven associated free energy changes: six binding energies and one folding energy (Fig. 4a, Extended Data Fig. 4a). As for RAF1 (Fig. 1k), the MoCHI binding energies for RALGDS correlate extremely well with independent *in vitro* measurements (Fig. 4b, c). The binding energies identify the known binding surfaces on KRAS, including the two known interfaces for SOS1 ^28^ (Fig. 2b, Extended Data Fig. 4b, median ROC-AUC=0.80, range=0.68-0.89 for weighted mean binding energies; median ROC-AUC=0.64, range=0.54-0.75 for weighted mean binding fitness measurements).

These seven free energy landscapes constitute >22,000 thermodynamic measurements, which is similar in scale to the number of measurements made for proteins in the entire scientific literature^29^.

### Specificity in the effector binding interface

We first considered how mutations in the binding interfaces alter binding to the six interaction partners. All six proteins bind KRAS through an overlapping set of contacts (Fig. 5a-c). This sharing of contacts is particularly pronounced for the three effector proteins, RAF1, PIK3CG and RALGDS (Fig. 5a). Comparing the mutational effects reveals that whereas some residues are critically important for binding to all three proteins, many substitutions alter the binding specificity (Fig. 5d). For example, many mutations in the negatively charged residues D33 and D38 and the hydrophobic residues I36 and Y40 strongly inhibit binding to all three effectors. However, a subset of hydrophobic substitutions at I36 inhibits binding to PIK3CG and RALGDS but not to RAF1 and substitution of L56 to negatively charged residues specifically increases binding to RAF1 whilst retaining binding to PIK3CG but inhibiting binding to RALGDS (Fig. 5d). In contrast, many substitutions at E37 inhibit binding to RAF1 and RALGDS but increase binding to PIK3CG. Mutating Y64 inhibits binding to PIK3CG and RALGDS but allows binding to RAF1. At S39 a subset of hydrophobic mutations inhibit binding to PIK3CG and RAF1 but not to RALGDS. Comparing the binding free energies for all six binding partners reveals a striking diversity of specificity changes that can be reached through single aa substitutions (Extended Data Fig. 5)

**Fig. 5.**
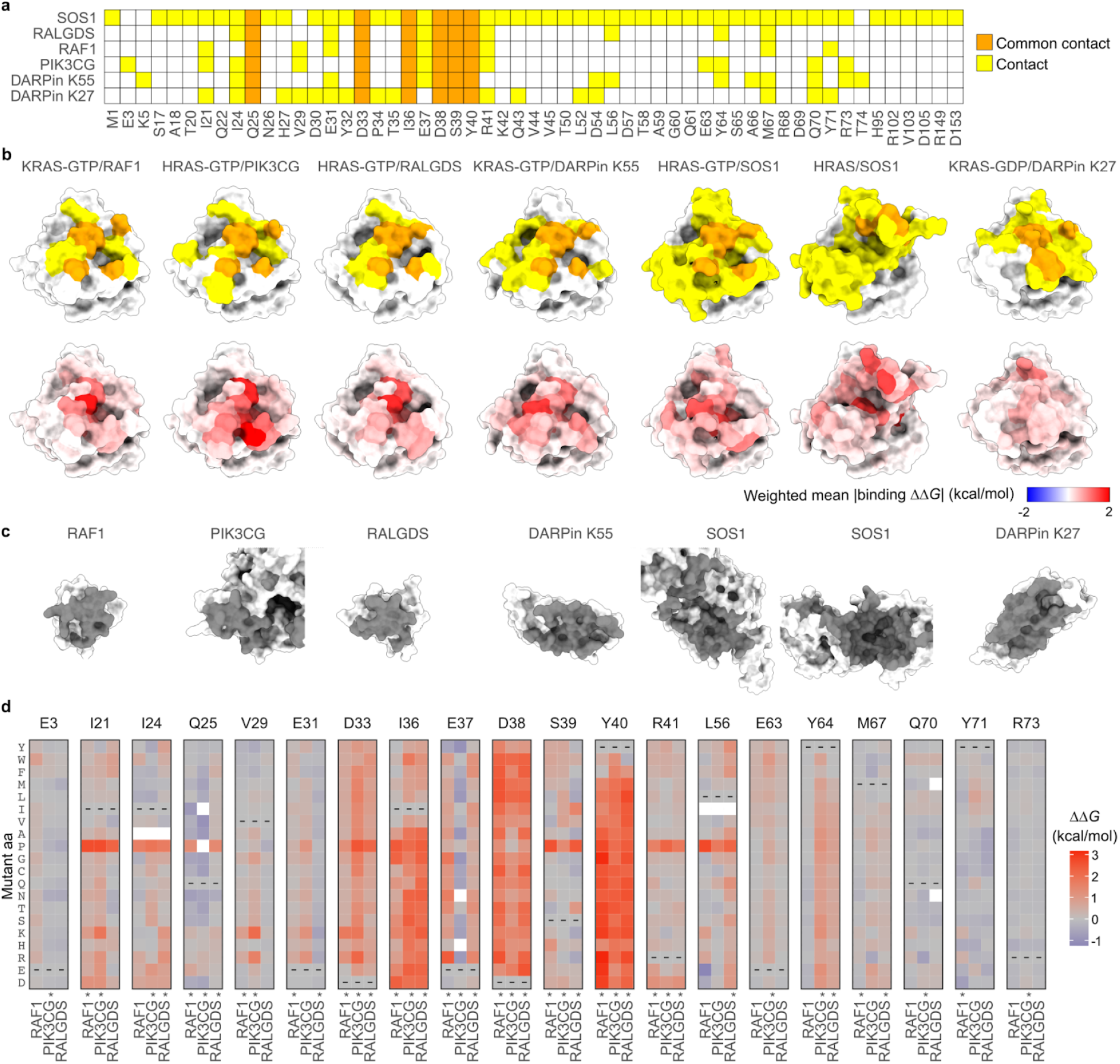
Energetic landscapes of KRAS interaction surfaces. **a**, Common and unique structural contacts between KRAS and the indicated six binding partners. **b**, 3D structures of KRAS indicating binding partner contacts (upper row, coloured as in **a**) and weighted mean absolute binding free energy change (lower row). **c**, 3D structures of binding partners (RAF1, 6VJJ; PIK3CG, 1HE8; RALGDS, 1LFD; DARPin K55, 5O2T; SOS1, 1NVW; DARPin K27, 5O2S) with binding interface indicated in grey. **d**, Heat maps of binding free energy changes in interface residues contacting at least one of the three effectors (RAF1, PIK3CG, RALGDS). Asterisks indicate binding interface residues for each partner.

### Inhibitory allosteric landscapes for six KRAS interactions

We next considered the specificity of mutational effects outside of the binding interfaces. We first focussed on the positions most enriched for allosteric mutations for each interaction, defining the major allosteric sites for each interaction as those in which the average absolute binding free energy change is as large or greater than the average across mutations in all the binding interfaces (Fig. 6a). Novel major allosteric sites were identified for all six binding partners, with a median of 9 major allosteric sites in the nucleotide-binding pocket and a median of 5.5 additional major allosteric sites for each interaction (Fig. 6a).

**Fig. 6.**
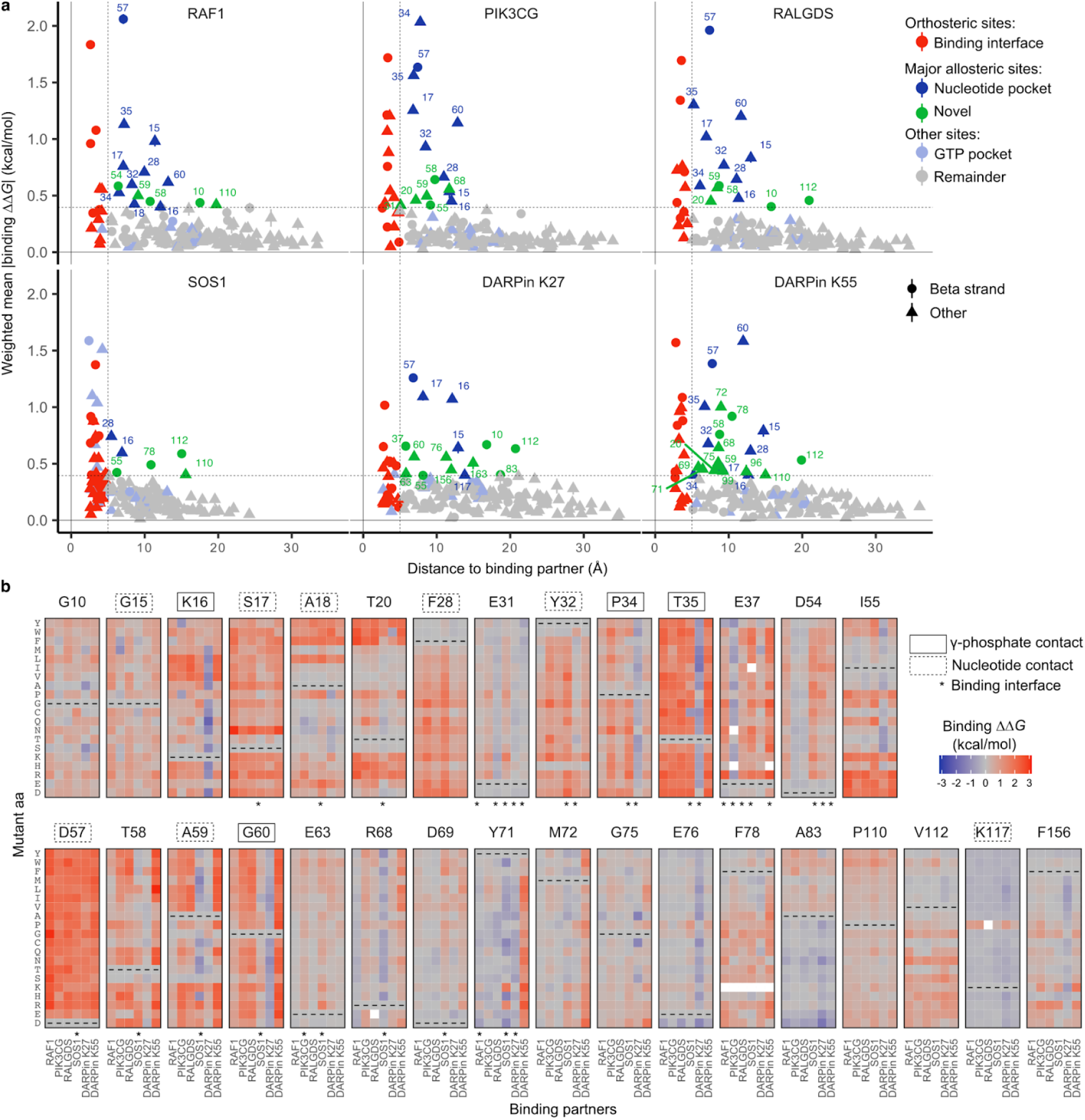
Allosteric control of binding specificity. **a**, Relationship between the weighted mean absolute change in free energy of binding and the distance to each corresponding binding partner (minimal side chain heavy atom distance). Major allosteric sites are defined as non-binding-interface residues with weighted mean absolute change in free energy of binding higher than the average of binding-interface residue mutations across all binding partners (horizontal dashed line). Error bars indicate 95% confidence interval (n ≥ 10)). **b**, Heat maps of binding free energy changes in all major allosteric sites. Nucleotide pocket and γ-phosphate-contacting residues are indicated.

We then compared the binding free energy changes between all six interaction partners for all mutations in these positions (Fig. 6b). Many substitutions at G10, G15, S17, D57, F78, P110 and V112 inhibit all 6 interactions (Fig. 6b). Substitutions of F28 to non-aromatic amino acids inhibit all 6 interactions, as do many changes to charged amino acids at I55 and to hydrophobic amino acids at A18 and A83 (Fig. 6b). Substitutions to P at I55, A59, R68, K117, F156 inhibit at least 5 interactions (Fig. 6b). Considering all mutations outside of the binding interface, allosteric mutations are enriched at G, P, F and T residues for 4/6 partners and depleted at charged residues for 6/6. Allosteric mutations are also enriched for substitutions to P for 6/6 partners and to R for 5/6 partners (Extended Data Fig. 7). The enrichment for allosteric mutations at G residues and for substitutions to P is also observed in two small protein domains ^12^.

### Allosteric control of binding specificity

That multiple mutations at many of the allosteric sites inhibit binding to all interaction partners suggests engagement of these sites is likely to generally inhibit KRAS function. However inspection of Figure 6b also reveals sets of mutations in the major allosteric sites that have more specific allosteric effects. Particularly striking examples are many mutations in residues K16, I55, G60 and F156 that allosterically inhibit binding to most KRAS interaction partners but allosterically increase binding to the DARPin K27 (Fig. 6b). DARPin K27 specifically recognises inactive GDP-bound KRAS and so mutations at these sites likely favour GDP-binding states. Consistent with this, K16 and G60 directly contact the γ-phosphate of GTP. Many substitutions of E76 also increase binding to DARPin K27 but with little effect on the other interactions. Additional examples include mutations at Y71 and M72 that specifically inhibit binding to DARPin K55 and mutations at D54 that inhibit four interactions but retain or enhance binding to PIK3CG and RALGDS (Fig. 6b). In addition, outside of these major allosteric sites there are many other mutations that allosterically alter both the binding affinity and specificity of KRAS (Extended Data Fig. 6).

## Discussion

We have presented here the first global map of inhibitory allosteric sites for any protein and the first comprehensive comparative map of the effects of mutations on the free energies of binding of a protein to multiple interaction partners. The dataset constitutes >22,000 free energy measurements, which is a rich resource for protein biophysics and computational biology.

KRAS is one of the most frequently mutated genes in cancer and one of the most sought after and valuable therapeutic targets. Our results reveal a number of principles concerning allosteric communication in KRAS. First, KRAS has many inhibitory allosteric sites. Second, most allosteric mutations inhibit binding to all three KRAS effectors, revealing the potential to broadly inhibit KRAS activity. Third, allosteric mutations are enriched close to binding sites, suggesting local energetic propagation as the main allosteric mechanism. Fourth, allosteric communication is anisotropic, with communication particularly effective across the central beta sheet of KRAS. Fifth, mutations can also allosterically control binding specificity, revealing extensive potential for regulatory, evolutionary and therapeutic modulation of signalling bias. Sixth, all four surface pockets of KRAS are allosterically active, with the effects of mutations in the distal and unexplored pocket 3 particularly striking. The comprehensive allosteric map therefore genetically validates all four pockets as suitable for therapeutic targeting and focuses attention on the largely ignored pocket 3.

The KRAS effector interface – like many protein surfaces – has to recognise structurally diverse proteins. Comprehensive mutagenesis of this surface shows that its evolution is constrained by fitness trade-offs, with mutations that increase binding to one protein typically having antagonistic pleiotropic effects on binding to others. However, the binding specificity of KRAS is highly evolvable, with single aa substitutions causing a diversity of specificity changes. These altered binding profiles can be useful experimental tools, providing ‘edgetic’ perturbations ^30^ to test the functions of individual molecular interactions and their combinations ^30,31^.

The accelerated pace of human genetics means we now know the proteins to therapeutically target in hundreds of human diseases ^32^. Unfortunately, however, effective therapeutics have only been developed against a small minority of these genetically-validated targets. In short, for many diseases we know the proteins to target but we do not know how to target them. For most proteins, we do not know the location of the ‘switches’ to target with drugs to turn them off or on. If we could find these switches, we would be able to develop drugs to control their activity.

The data presented here and in additional recent studies ^12,33–36^ have revealed that allosteric sites are much more prevalent than is widely appreciated. Moreover, the approach that we have applied here to KRAS is quite general and can be used to identify allosteric sites in many different proteins. We believe that using this general strategy it will be possible to systematically map the regulatory sites to target in many important proteins. Mapping allosteric sites is likely to play an increasingly important role in drug development, laying the foundations for therapeutically targeting proteins previously considered to be ‘undruggable’.

## Methods

### Media and buffers

- LB: 10 g/L Bacto-tryptone, 5 g/L Yeast extract, 10 g/L NaCl. Autoclaved 20 min at 120ºC.
- YPDA: 20 g/L glucose, 20 g/L Peptone, 10 g/L Yeast extract, 40 mg/L adenine sulphate. Autoclaved 20 min at 120ºC.
- SORB: 1 M sorbitol, 100 mM LiOAc, 10 mM Tris pH 8.0, 1 mM EDTA. Filter sterilized (0.2 mm Nylon membrane, ThermoScientific).
- Plate mixture: 40% PEG3350, 100 mM LiOAc, 10 mM Tris-HCl pH 8.0, 1 mM EDTA pH 8.0. Filter sterilised.
- Recovery medium: YPD (20 g/L glucose, 20 g/L Peptone, 10 g/L Yeast extract) + 0.5 M sorbitol. Filter sterilised.
- SC -URA: 6.7 g/L Yeast Nitrogen base without amino acid, 20 g/L glucose, 0.77 g/L complete supplement mixture drop-out without uracil. Filter sterilised.
- SC -URA/MET/ADE: 6.7 g/L Yeast Nitrogen base without amino acid, 20 g/L glucose, 0.74 g/L complete supplement mixture drop-out without uracil, adenine and methionine. Filter sterilised.
- Competition medium: SC –URA/MET/ADE + 200 ug/mL methotrexate (BioShop Canada Inc., Canada), 2% DMSO.
- DNA extraction buffer: 2% Triton-X, 1% SDS, 100mM NaCl, 10mM Tris-HCl pH8, 1mM EDTA pH8.

### Plasmid construction

Two generic plasmids were constructed to be able to assay any protein of interest by BindingPCA or AbundancePCA: the BindingPCA plasmid (pGJJ161) and the AbundancePCA plasmid (pGJJ162).

The BindingPCA plasmid (pGJJ161) and AbundancePCA plasmid (pGJJ162) were derived from the previous BindingPCA plasmid (pGJJ001) and the previous AbundancePCA plasmid (pGJJ045) ^12^. The C-terminus (GGGGS)4 linker of DHRF3 were changed to N-terminus which allowed us to fuse protein of interest to N-terminus to the DHFR3 fragment in both abundance and binding PCA assays.

One KRAS AbundancePCA plasmid, 6 BindingPCA plasmids and one KRAS mutagenesis plasmid are used in this paper. To construct the KRAS AbundancePCA plasmid (pGJJ271), the sequence of full length KRAS (188 aa) was amplified from a plasmid, a gift from Luis Serrano lab using primer pair oGJJ231/oGJJ232 (Supplementary Table 1). This primer pair also introduced the HindIII and NheI restriction sites. The PCR product was digested by HindIII and NheI then was cloned into the digested pGJJ162 plasmid using T4 Ligase (NEB). To construct 6 KRAS BindingPCA plasmids, a common KRAS bindingPCA plasmid (pGJJ317) was constructed by ligating full length KRAS sequence digested by HindIII and NheI to digested BindingPCA plasmid. 6 BindingPCA plasmids are constructed by ligating each binding partners PCR product which was digested by BamHI and SpeI to digested pGJJ317 using T4 Ligase (NEB). To construct RAF1 bindingPCA plasmid (pGJJ336), the sequence of RAF1RBD (52-131) was amplified from the cDNA of 293T cell line using primer pair oGJJ74/oGJJ307 which also introduced the BamHI and SpeI restriction sites. To construct PI3KCG bindingPCA plasmid (pGJJ565), the sequence of PIK3CG RBD (203-312) was amplified from R777-E169 Hs.PIK3CG (addgene) using primer pair oWCC169/oWCC170. To construct RALGDS bindingPCA plasmid (pGJJ400), the sequence of RALGDS RBD (778-864) was amplified from R777-E169 Hs.PIK3CG (addgene) using primer pair oWCC28/oWCC29. To construct SOS1 bindingPCA plasmid (pGJJ541), the sequence of SOS1 (564-1049) was amplified from plasmid R777-E317 Hs.SOS1 (addgene) using primer pair oWCC149/oWCC150. To construct DARPin K27 bindingPCA plasmid (pGJJ553), the sequence of DARPin K27 was amplified from plasmid pCASP-SptP120-K27-HilA (addgene) using primer pair oWCC157/oWCC158. To construct DARPin K55 bindingPCA plasmid (pGJJ554), the sequence of DARPin K55 was amplified from plasmid pCASP-SptP120-K55-HilA (addgene) using primer pair oWCC159/oWCC160. To construct the KRAS mutagenesis plasmid (pGJJ380), pGJJ191 plasmid was constructed firstly which contained a streptomycin resistance gene cassette. The pGJJ191 plasmid was amplified in two fragment, one ori cassette which also contained AvrII and HindIII restriction sites using primer pair oGJJ308/oGJJ309, the other streptomycin resistance gene cassette using primer pair oGJJ310/oGJJ311, which were then assembled by Gibson reaction (prepared in house) at 50ºC for one hour. KRAS was digested by AvrII and HindIII from abundancePCA plasmid and ligated into digested pGJJ191. Then a BbvCI restriction site was introduced using primer pair oWCC51/oWCC52.

### Mutagenesis library construction

The plasmid-based one-pot saturation (nicking) mutagenesis protocol was used in this study ^13^. KRAS are divided to three blocks in order to be fully sequenced by Illumina paired-end 150 NextSeq pipeline.

An initial single round of nicking mutagenesis using equimolar mixes of degenerate KRAS primers (Supplementary Table 3) was obtained for two reasons: (1) To obtain random single mutants to use as template for another round of nicking mutagenesis (by randomly selecting single colonies and verified by Sanger sequencing) and (2) to quantify the degenerate primer positional bias and compensate for it in the shallow double mutant libraries.

To construct three final KRAS libraries, an equimolar pool of single mutants of each block and wild type were used as the plasmid template for a round of nicking mutagenesis. To compensate for the extreme positional biases, each mutagenic primer was mixed in the pool inversely to the mean read counts per position from these first-round nicking libraries.

The libraries midi-preps were digested with HindIII and NheI restriction enzymes and the insert containing the mutated protein was gel purified (MinElute Gel Extraction Kit, QIAGEN) to be later cloned into the AbundancePCA plasmid and BindingPCA plasmids by temperature-cycle ligation. The AbundancePCA plasmid and BindingPCA plasmids were all digested by HindIII and NheI enzymes and purified using the QIAquick Gel Extraction Kit (QIAGEN). The assembly of AbundancePCA libraries and BindingPCA libraries were done overnight by temperature-cycle ligation using T4 ligase (New England Biolabs) according to the manufacturer’s protocol, 67 fmol of backbone and 200 fmol of insert in a 33.3 uL reaction. The ligation was desalted by dialysis using membrane filters for 1h and later concentrated 3.3X using a SpeedVac concentrator (Thermo Scientific).

All concentrated assembled libraries were transformed into NEB 10β High-efficiency Electrocompetent E. coli cells according to the manufacturer’s protocol (volumes used in each library specified in Supplementary Table 2). Cells were allowed to recover in SOC medium (NEB 10β Stable Outgrowth Medium) for 30 minutes and later transferred to 200 mL of LB medium with ampicillin 4X overnight. The total number of estimated transformants for each library can be found in Supplementary Table 2. 100 mL of each saturated E. coli culture were harvested next morning to extract the plasmid library using the QIAfilter Plasmid Midi Kit (QIAGEN).

### Methotrexate selection assays

The methotrexate selection assay protocol was described in our previous study^12^. The high-efficiency yeast transformation protocol was scaled in volume depending on the targeted number of transformants of each library. The transformation protocol described below (adjusted to a pre-culture of 175 mL of YPDA) was scaled up or down in volume as reported in Supplementary Table 2.

For each of the selection assays (3 blocks x 6 BindingPCA + 3 blocks x 1 AbundancePCA), three independent pre-cultures of BY4742 were grown in 20 mL standard YPDA at 30ºC overnight. The next morning, the cultures were diluted into 175 mL of pre-wormed YPDA at an OD600nm = 0.3. The cultures were incubated at 30ºC for 4 hours. After growth, the cells were harvested and centrifuged for 5 minutes at 3,000g, washed with sterile water and later with SORB medium (100mM LiOAc, 10mM Tris pH 8.0, 1mM EDTA, 1M sorbitol). The cells were resuspended in 8.6 mL of SORB and incubated at room temperature for 30 minutes. After incubation, 175 μL of 10mg/mL boiled salmon sperm DNA (Agilent Genomics) was added to each tube of cells, as well as 3.5 μg of plasmid library. After gentle mixing, 35 mL of Plate Mixture (100mM LiOAc, 10mM Tris-HCl pH 8, 1mM EDTA/NaOH, pH 8, 40% PEG3350) were added to each tube to be incubated at room temperature for 30 more minutes. 3.5 mL of DMSO was added to each tube and the cells were then heat shocked at 42ºC for 20 minutes (inverting tubes from time to time to ensure homogenous heat transfer). After heat shock, cells were centrifuged and re-suspended in ∼50 mL of recovery media and allowed to recover for 1 hour at 30ºC. Next, cells were again centrifuged, washed with

SC-URA medium and re-suspended in SC -URA (volume used in each library found in Supplementary Table 2). After homogenization by stirring, 10 uL were plated on SC -URA Petri dishes and incubated for ∼48 hours at 30ºC to measure the transformation efficiency. The independent liquid cultures were grown at 30ºC for ∼48 hours until saturation. The number of yeast transformants obtained in each library assay can be found in Supplementary Table 2.

For each of the BindingPCA or AbundancePCA assays, each of the growth competitions was performed right after yeast transformation. After the first cycle of post-transformation plasmid selection, a second plasmid selection cycle (input) was performed by inoculating SC -URA/MET/ADE at a starting OD600nm = 0.1 with the saturated culture (volume of each experiment specified in Supplementary Table 2). Cells were grown for 4 generations at 30ºC under constant agitation at 200 rpm. This allowed the pool of mutants to be amplified and enter the exponential growth phase. The competition cycle (output) was then started by inoculating cells from the input cycle into the competition media (SC -URA/MET/ADE + 200 ug/mL Methotrexate) so that the starting OD600nm was 0.05. For that, the adequate volume of cells was collected, centrifuged at 3,000 rpm for 5 minutes and resuspended in the pre-warmed output media. Meanwhile, each input replicate culture was splitted in two and harvested by centrifugation for 5 min at 5,000g at 4ºC. Yeast cells were washed with water, pelleted and stored at -20ºC for later DNA extraction. After ∼5 generations of competition cycle, each output replicate culture was splitted into two and harvested by centrifugation for 5 min at 5,000g at 4ºC, washed twice with water and pelleted to be stored at -20ºC.

### DNA extractions and plasmid quantification

The DNA extraction protocol used was described in our previous study^12^. A 50 mL harvested culture of OD600nm ∼ 1.6 is described below. Cell pellets (one for each experiment input/output replicate) were re-suspended in 1 mL of DNA extraction buffer, frozen by dry ice-ethanol bath and incubated at 62ºC water bath twice. Subsequently, 1 mL of Phenol/Chloro/Isoamyl 25:24:1 (equilibrated in 10mM Tris-HCl, 1mM EDTA, pH8) was added, together with 1 g of acid-washed glass beads (Sigma Aldrich) and the samples were vortexed for 10 minutes. Samples were centrifuged at RT for 30 minutes at 4,000 rpm and the aqueous phase was transferred into new tubes. The same step was repeated twice. 0.1 mL of NaOAc 3M and 2.2 mL of pre-chilled absolute ethanol were added to the aqueous phase. The samples were gently mixed and incubated at -20ºC for 30 minutes. After that, they were centrifuged for 30 min at full speed at 4ºC to precipitate the DNA. The ethanol was removed and the DNA pellet was allowed to dry overnight at RT. DNA pellets were resuspended in 0.6 mL TE 1X and treated with 5 uL of RNaseA (10mg/mL, Thermo Scientific) for 30 minutes at 37ºC. To desalt and concentrate the DNA solutions, QIAEX II Gel Extraction Kit was used (50 µL of QIAEX II beads). The samples were washed twice with PE buffer and eluted twice by 125 µL of 10 mM Tris-HCI buffer, pH 8.5 and then combined two elution. Finally, plasmid concentrations in the total DNA extract (that also contained yeast genomic DNA) were quantified by qPCR using the primer pair oGJJ152-oGJJ153, that binds to the ori region of the plasmids.

### Sequencing library preparation

The sequencing library preparation protocol was described in our previous study^12^. The sequencing libraries were constructed in two consecutive PCR reactions. The first PCR (PCR1) was designed to amplify the mutated protein of interest and to increase the nucleotide complexity of the first sequenced bases by introducing frame-shift bases between the adapters and the sequencing region of interest. The second PCR (PCR2) was necessary to add the remainder of the Illumina adapter and demultiplexing indexes.

To avoid PCR biases, PCR1 of each independent sample (input/output replicates of any of the yeast assays) was run with an excess of plasmid template 20-50 times higher than the number of expected sequencing reads per sample. Each reaction started with a maximum of 1.25×10^7^ template plasmid molecules per uL of PCR1, avoiding introducing more yeast genomic DNA that interfered with the efficiency of the PCR reaction. For this reason, PCR1s were scaled up in volume as specified in Supplementary Table 2. The PCR1 reactions were run using Q5 Hot Start High-Fidelity DNA Polymerase (New England Biolabs) according to the manufacturer’s protocol, with 25 pmol of pooled frame-shift primers as specified in Supplementary Table 1 for difference blocks (forward and reverse primers were independently pooled according to the nucleotide diversity of each oligo, Supplementary Table 1). The PCR reactions were set to 60ºC annealing temperature, 10 seconds of extension time and run for 15 cycles. Excess primers were removed by adding 0.04 uL of ExoSAP-IT (Affymetrix) per uL of PCR1 reaction and incubated for 20 min at 37 ̊C followed by an inactivation for 15 min at 80 ̊C. The PCRs of each sample were then pooled and purified using the MinElute PCR Purification Kit (QIAGEN) according to the manufacturer’s protocol. DNA was eluted in EB to a volume 6 times lower than the total volume of PCR1.

PCR2 reactions were run for each sample independently using Hot Start High-Fidelity DNA Polymerase. The total reaction of PCR2 was reduced to half of PCR1, using 0.05 uL of the previous purified PCR1 per uL of PCR2. In this second PCR the remaining parts of the Illumina adapters were added to the library amplicon. The forward primer (5’ P5 Illumina adapter) was the same for all samples, while the reverse primer (3’ P7 Illumina adapter) differed by the barcode index (oligo sequences in Supplementary Table 1), to be subsequently pooled together and demultiplex them after deep sequencing (indexes used in each replicate of each sequencing run found in Supplementary Table 2). 8 cycles of PCR2s were run at 62ºC of annealing temperature and 10 seconds of extension time. All reactions from the same sample were pooled together and an aliquot was run on a 2% agarose gel to be quantified. All samples were purified using the QIAEX II Gel Extraction Kit. The purified amplicon library pools were subjected to Illumina 150bp paired-end NextSeq sequencing at the CRG Genomics Core Facility.

### Sequencing data processing

FastQ files from paired-end sequencing of all BindingPCA and AbundancePCA experiments were processed with DiMSum v1.2.9 ^38^ using default settings with minor adjustments: https://github.com/lehner-lab/DiMSum. Supplementary Table 4 contains DiMSum fitness estimates and associated errors for all experiments. Experimental design files and command-line options required for running DiMSum on these datasets are available on GitHub (https://github.com/lehner-lab/krasddpcams). In all cases, adaptive minimum Input read count thresholds based on the corresponding number of nucleotide substitutions (“fitnessMinInputCountAny” option) were selected in order to minimise the fraction of reads per variant related to sequencing error-induced “variant flow” from lower order mutants.

Variant counts associated with all samples (output from DiMSum stage 4) were further filtered using a custom script to retain only those variants with single aa substitutions including a G/T in the third codon position (encoded by “NNK”) or aa substitutions representing high confidence backgrounds. The latter were defined as single aa substitutions observed at least 200 times (in different double aa variants) in at least five (out of a total of seven) *BindingPCA*/*AbundancePCA* experiments. For double aa variants, we required one of the constituent single aa variants to be a high confidence background mutation. All read counts associated with remaining single or double aa variants (likely the result of PCR and sequencing errors) were discarded. Finally, fitness estimates and associated errors were then obtained from the resulting filtered variant counts with DiMSum (“countPath” option).

### Thermodynamic model fitting with MoCHI

We used MoCHI (https://github.com/lehner-lab/MoCHI) to fit a global mechanistic model to all 21 ddPCA datasets (7 phenotypes x 3 blocks) simultaneously. The software is based on our previously described genotype-phenotype modelling approach ^12^ with additional functionality and improvements for ease-of-use and flexibility.

Briefly, we model individual KRAS PPIs as an equilibrium between three states: unfolded and unbound (*uu*), folded and unbound (*fu*), and folded and bound (*fb*). We assume that the probability of the unfolded and bound state (*ub*) is negligible and free energies of folding and binding are additive i.e. the total binding and folding free energy changes of an arbitrary variant relative to the wild-type sequence is simply the sum over residue-specific energies corresponding to all constituent single amino acid substitutions. Furthermore, we assume binding energies are specific for each binding partner whereas folding energies are shared/intrinsic to KRAS i.e. unaffected by the identity/presence/expression of a given binding partner.

We configured MoCHI parameters to specify a neural network architecture consisting of seven additive trait layers (free energies) i.e. one for each biophysical trait to be inferred (6x binding and 1x folding), as well as one linear transformation layer per experiment (3x *abundancePCA* and 18x *bindingPCA* fitness). The specified non-linear transformations “TwoStateFractionFolded” and “ThreeStateFractionBound” derived from the Boltzmann distribution function relate energies to proportions of folded and bound molecules respectively. The target (output) data to fit the neural network comprises fitness scores for wild-type, single and double aa substitution variants from all 21 ddPCA datasets.

A random 30% of double aa substitution variants was held out during model training, with 20% representing the validation data and 10% representing the test data. Validation data was used to evaluate training progress and optimise hyperparameters (batch size). Optimal hyperparameters were defined as those resulting in the smallest validation loss after 100 training epochs. Test data was used to assess final model performance.

MoCHI optimises the parameters θ of the neural network using stochastic gradient descent on a loss function ℒ [θ] based on a weighted and regularised form of mean absolute error:

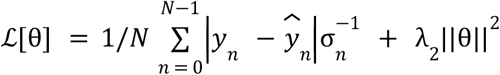

where *y*_*n*_ and *σ*_*n*_ are the observed fitness score and associated standard error respectively for variant *n, ŷ*_*n*_ is the predicted fitness score, *N* is the batch size and *λ*_2_ is the *L*_2_ regularisation penalty. In order to penalise very large free energy changes (typically associated with extreme fitness scores) we set *λ*_2_ to 10^−6^ representing light regularisation. The mean absolute error is weighted by the inverse of the fitness error 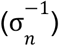 in order to downweight the contribution of less confidently estimated fitness scores to the loss. Furthermore, in order to capture the uncertainty in ddPCA fitness estimates, the training data was replaced with a random sample from the fitness error distribution of each variant. The validation and test data was left unaltered.

Models were trained with default settings i.e. for a maximum of 1000 epochs using the Adam optimization algorithm with an initial learning rate of 0.05. MoCHI reduces the learning rate exponentially (γ = 0.98) if the validation loss has not improved in the most recent ten epochs compared to the preceding ten epochs. In addition, MoCHI stops model training early if the wild-type free energy terms over the most recent ten epochs have stabilised (standard deviation ≤ 10^−3^).

Free energies are calculated directly from model parameters as follows: Δ*G*_*b*_ = θ_*b*_*RT* and Δ*G*_*f*_ = θ_*f*_*RT*, where *T* = 303 K and *R* = 0.001987 kcalK^-1^mol^-1^. We estimated the confidence intervals of model-inferred free energies using a Monte Carlo simulation approach. The variability of inferred free energy changes was calculated between ten separate models fit using data from [1] independent random training-validation-test splits and [2] independent random samples of fitness estimates from their underlying error distributions. Confident inferred free energy changes are defined as those with Monte Carlo simulation derived 95% confidence intervals < 1 kcal/mol. Supplementary Table 5 contains inferred binding and folding free energy changes of mutations for all binding partners.

## Acknowledgements

This work was funded by European Research Council (ERC) Advanced (883742) and Consolidator (616434) grants, the Spanish Ministry of Science and Innovation (BFU2017-89488-P, EMBL Partnership, Severo Ochoa Centre of Excellence), the Bettencourt Schueller Foundation, the AXA Research Fund, Agencia de Gestio d’Ajuts Universitaris i de Recerca (AGAUR, 2017 SGR 1322), and the CERCA Program/Generalitat de Catalunya. C.W. was funded by an EMBO long-term fellowship (ALTF 881-2020). We thank all members of the Lehner Lab for helpful discussions and suggestions.

## Author contributions

B.L. and C.W. conceived the project and designed the experiments; C.W. performed the experiments; C.W. and A.J.F. performed the data analysis; B.L., C.W., and A.J.F. wrote the manuscript.

## Additional information

Supplementary Information is available for this paper. Correspondence and requests for materials should be addressed to B.L.

## Data availability

All DNA sequencing data have been deposited in the Sequence Read Archive (SRA) under BioProject PRJNA907205: https://www.ncbi.nlm.nih.gov/bioproject/PRJNA907205. All fitness measurements and free energies are provided in Supplementary tables 4 and 5.

## Code availability

Source code for fitting thermodynamic models (MoCHI) is available at https://github.com/lehner-lab/MoCHI. Source code for all downstream analyses and to reproduce all figures described here is available at https://github.com/lehner-lab/krasddpcams.

## Extended Data

**Extended Data Fig. 1.**
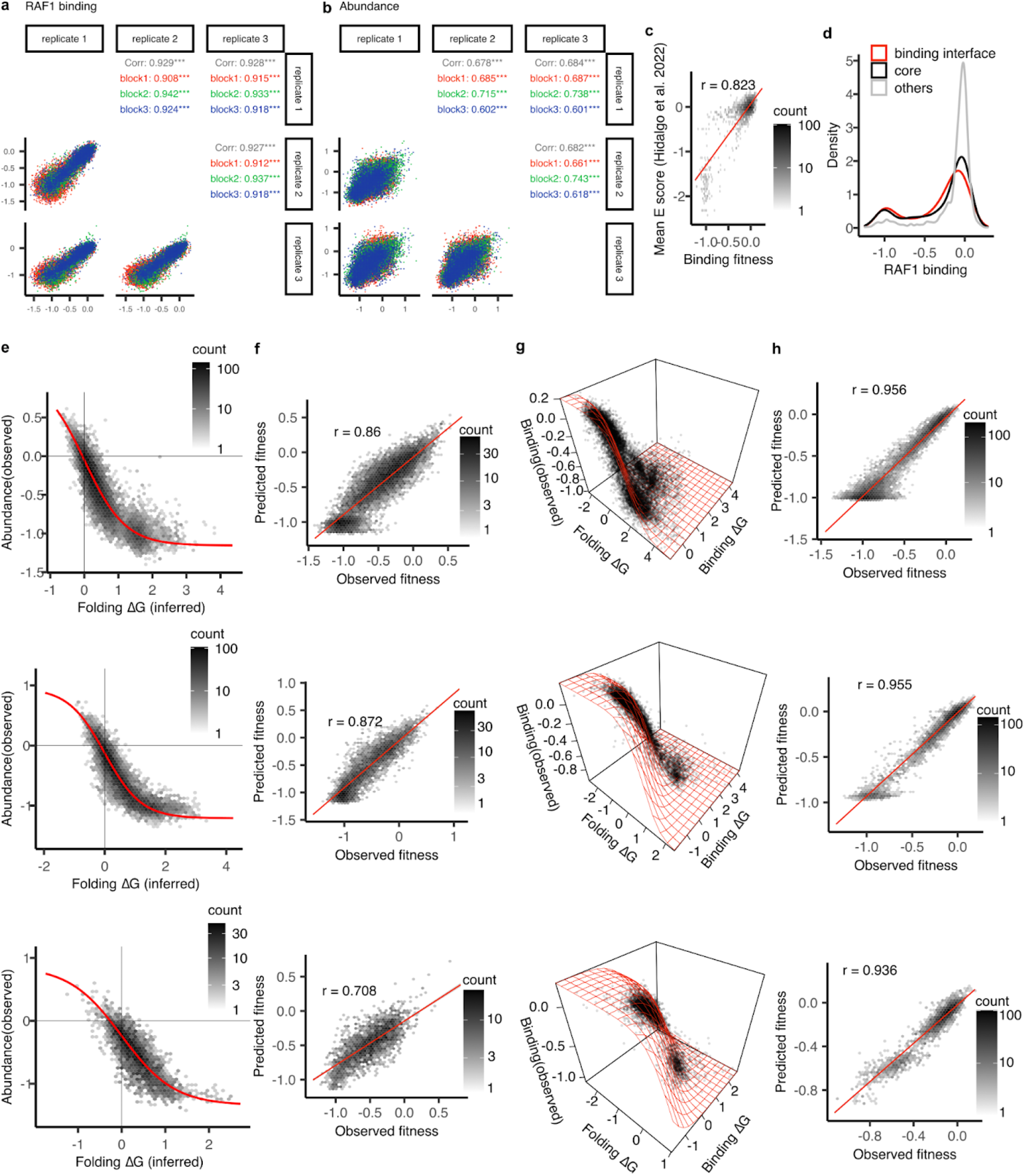
Experimental reproducibility and thermodynamic model fitting. **a**,**b**, Scatter plots showing the reproducibility of each block’s binding (a) and abundance (b) fitness estimates from ddPCA. Pearson’s r indicated on the top right corner. **c**, Comparison of the binding fitness to previously reported KRAS-RAF1 binding E score^17^. Pearson’s r = 0.82. **d**, single mutation fitness density distributions. **e**, Non-linear relationships (global epistasis) between observed AbundancePCA fitness and changes in free energy of folding. **f**, Scatter plots of predicted abundance fitness against observed abundance fitness from the model. **g**, Non-linear relationships (global epistasis) between observed. BindingPCA fitness and both free energies of binding and folding. **h**, Predicted binding fitness against observed binding fitness. Three rows stand for three mutagenesis library blocks (block 1, on the top, block 2, in the middle, block 3, on the bottom).

**Extended Data Fig. 2.**
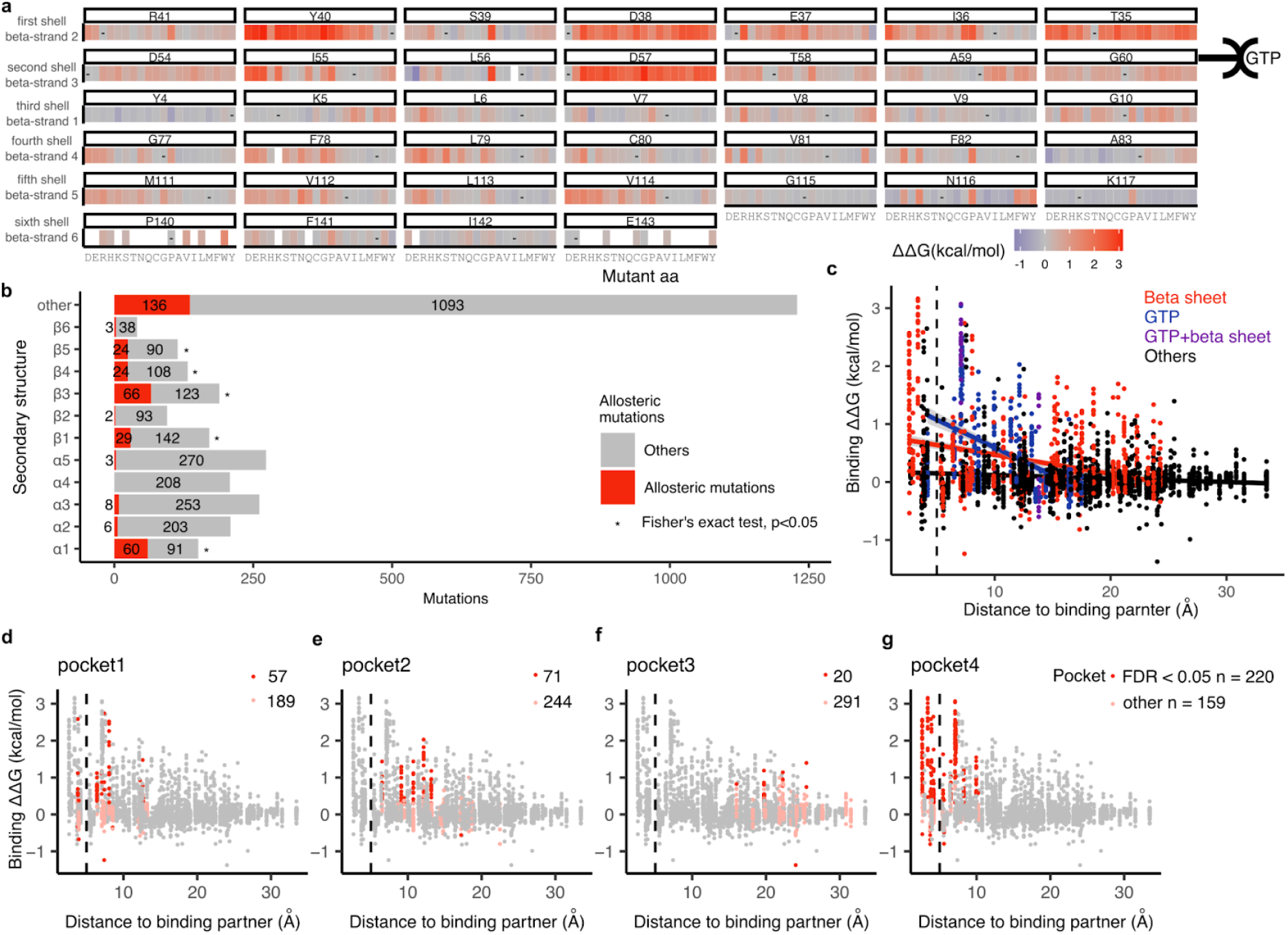
Allosteric mutations in the KRAS beta sheet and surface pockets. **a**, Heat maps of binding free energy changes of residues in the beta sheet. GTP indicates the location of GTP in the 3D structure. **b**, Number of allosteric mutations in each secondary structure element. *, odds ratio > 1, and Fisher’s exact two sided test, p < 0.05. **c**, Scatter plot showing the binding free energy changes of all mutations and the distance to the binding partner. Residues in beta sheet and GTP binding sites (minimal side chain heavy atom distance to GTP < 5 Å) are coloured as indicated. **d, e, f, g**, Scatter plot showing the binding free energy changes of all mutations and the distance to the binding partner. Residues in each pocket are coloured as indicated.

**Extended Data Fig. 3.**
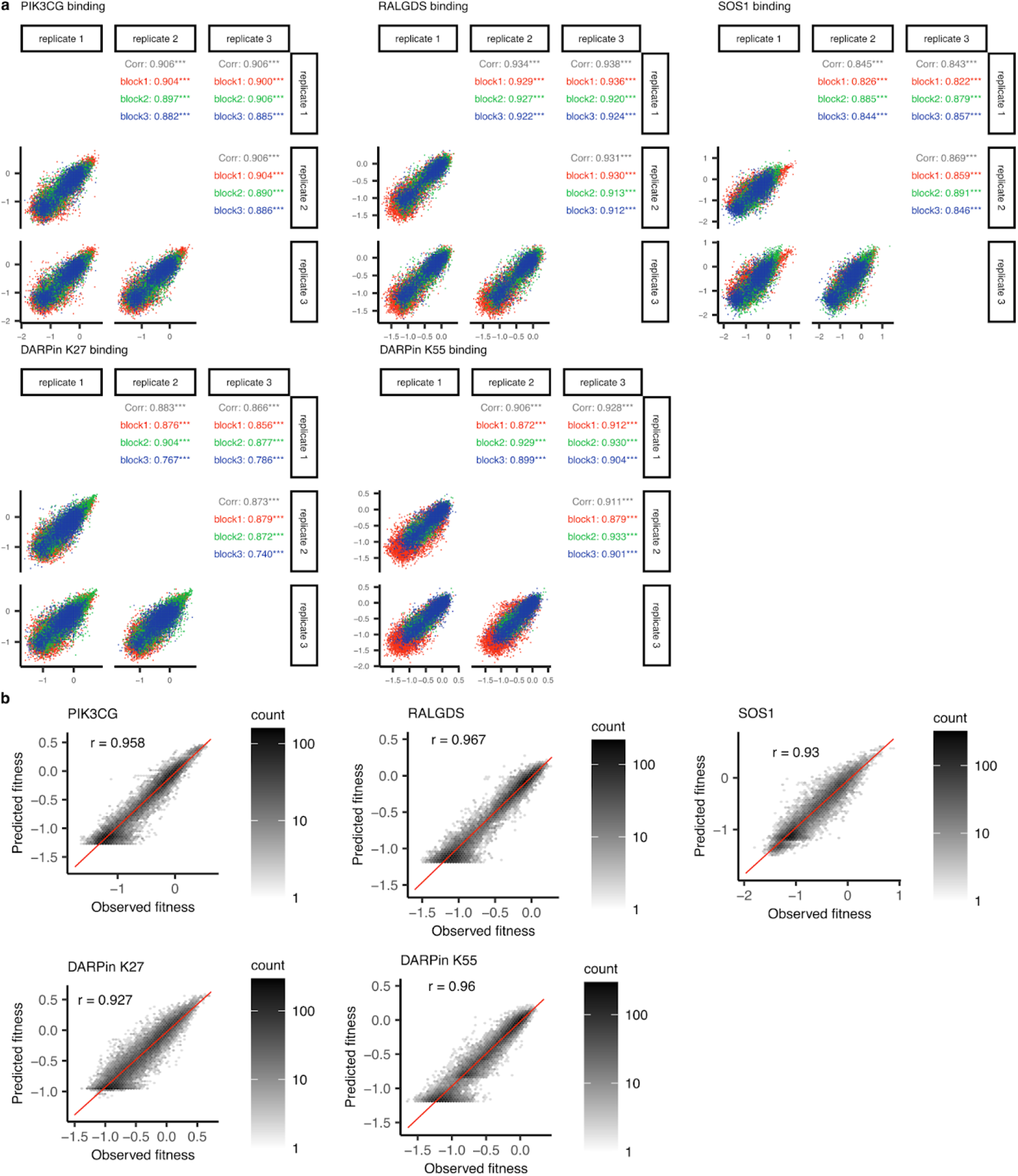
Experimental reproducibility and thermodynamic model fitting for five additional interaction partners. **a**, Scatter plots showing the reproducibility of each block’s binding fitness estimates from ddPCA. Pearson’s r indicated on the top right corner. **b**, Performance of models fit to ddPCA data. Scatter plots of predicted binding fitness against observed binding fitness.

**Extended Data Fig. 4.**
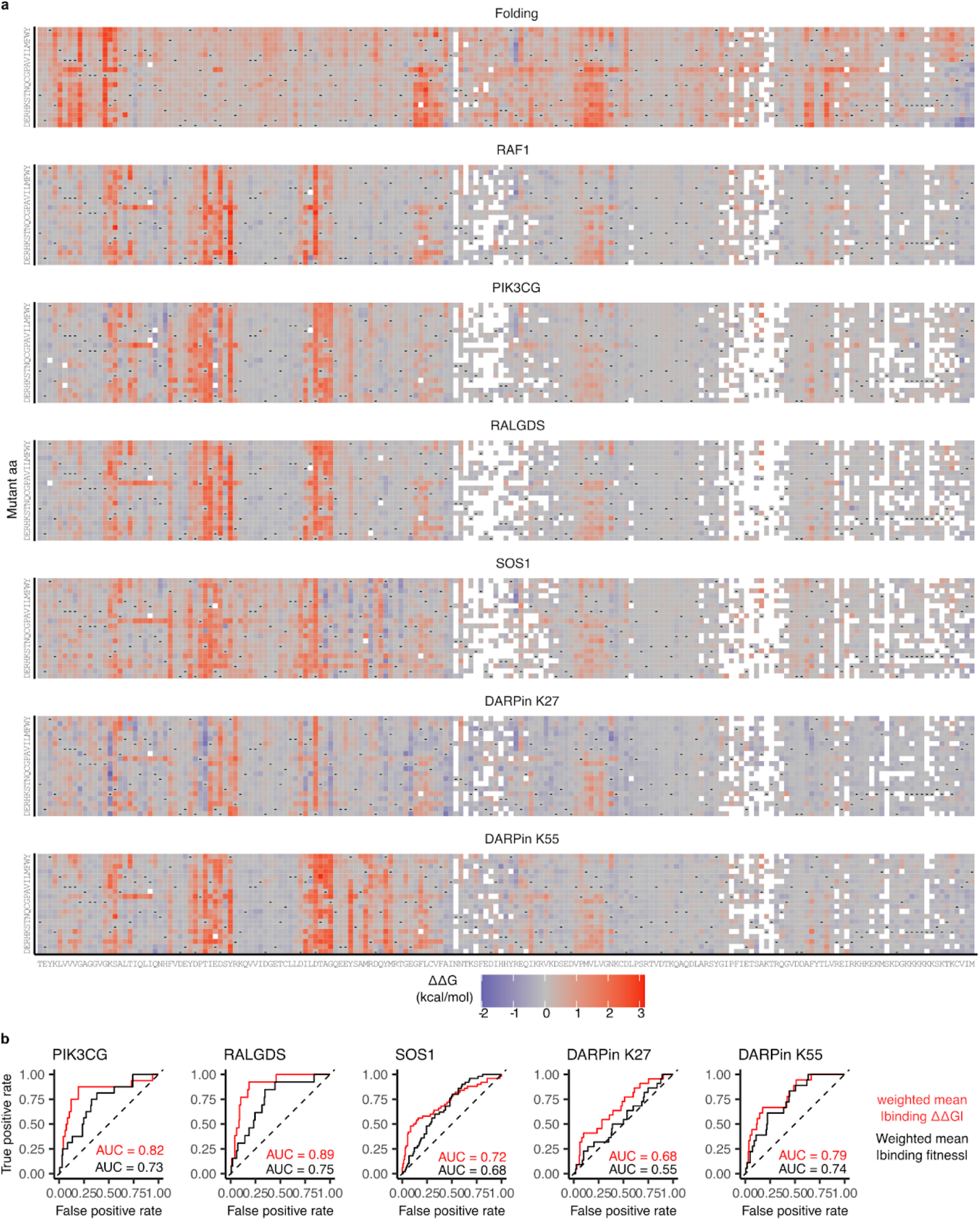
Seven KRAS free energy landscapes. **a**, Heat maps of folding and binding free energy changes. **b**, ROC curves for predicting binding interface residues (distance to binding partner < 5 Å) using weighted mean absolute binding free energy changes (ΔΔG) in red or using weighted mean absolute binding fitness in black. AUC = Area Under the Curve.

**Extended Data Fig. 5.**
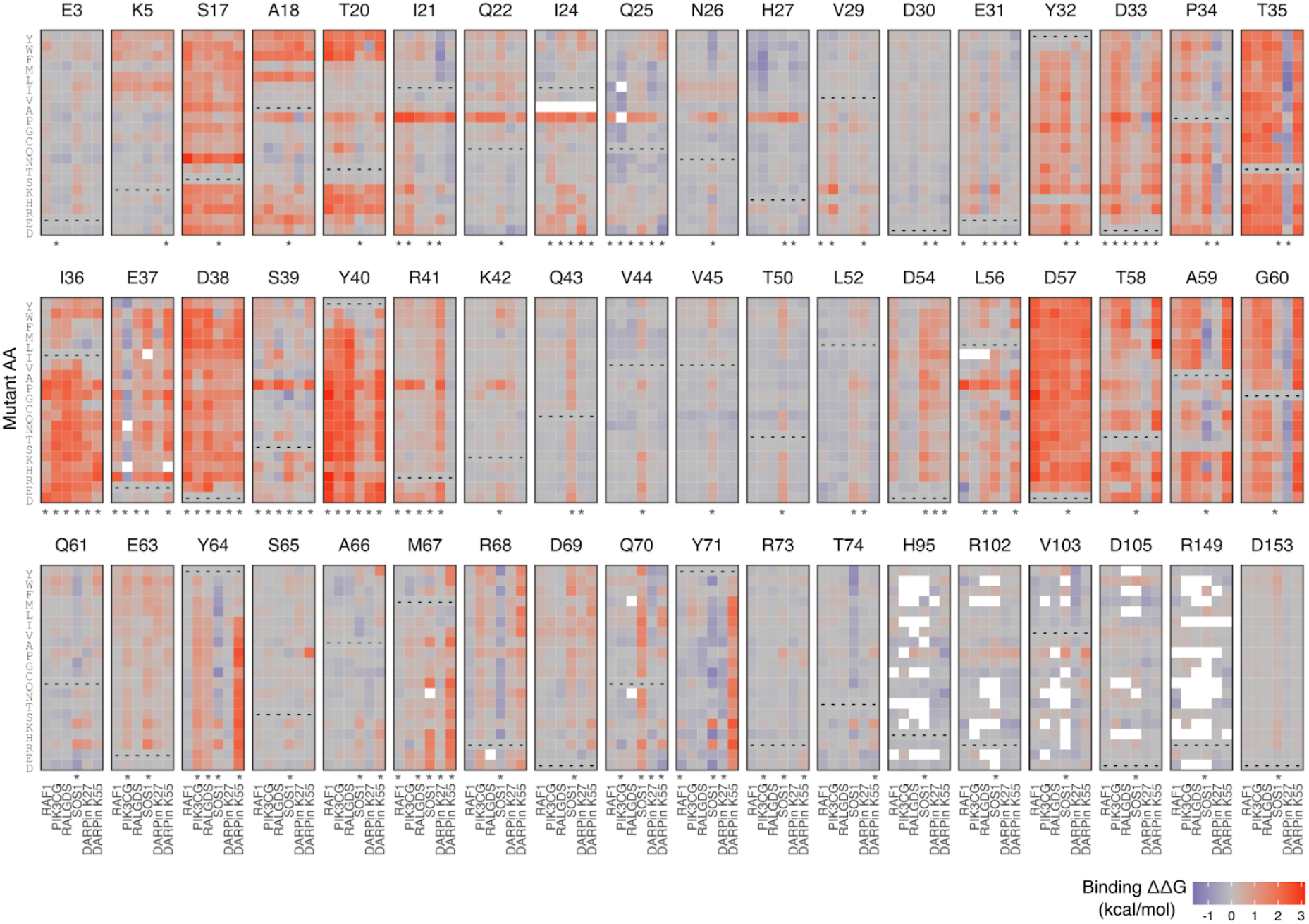
Binding interface specificity for all interactions. Heat maps of binding free energy changes of all binding partners (RAF1, PIK3CG, RALGDS, SOS1, DARPin K27, DARPin K55) binding interface residues. *, binding interface residues of each binding partner.

**Extended Data Fig. 6.**
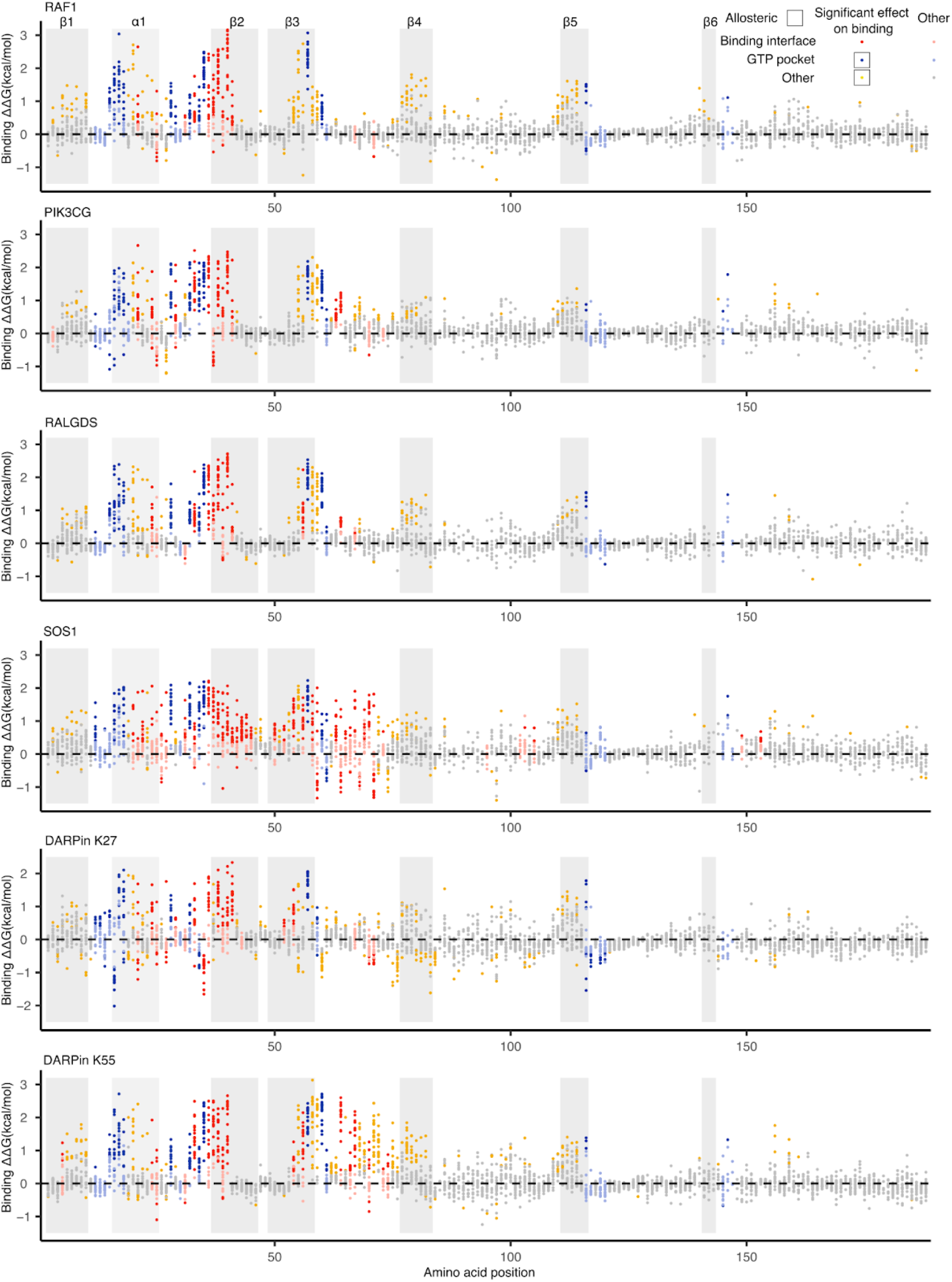
Binding energy and allosteric landscapes for all six binding partners. Scatter plot showing the binding free energy changes of all mutations coloured according to residue position and whether the free energy change is larger than the weighted mean of binding free energy changes in the binding-interfaces of all six proteins. Two-sided Z test, FDR < 0.05.

**Extended Data Fig. 7.**
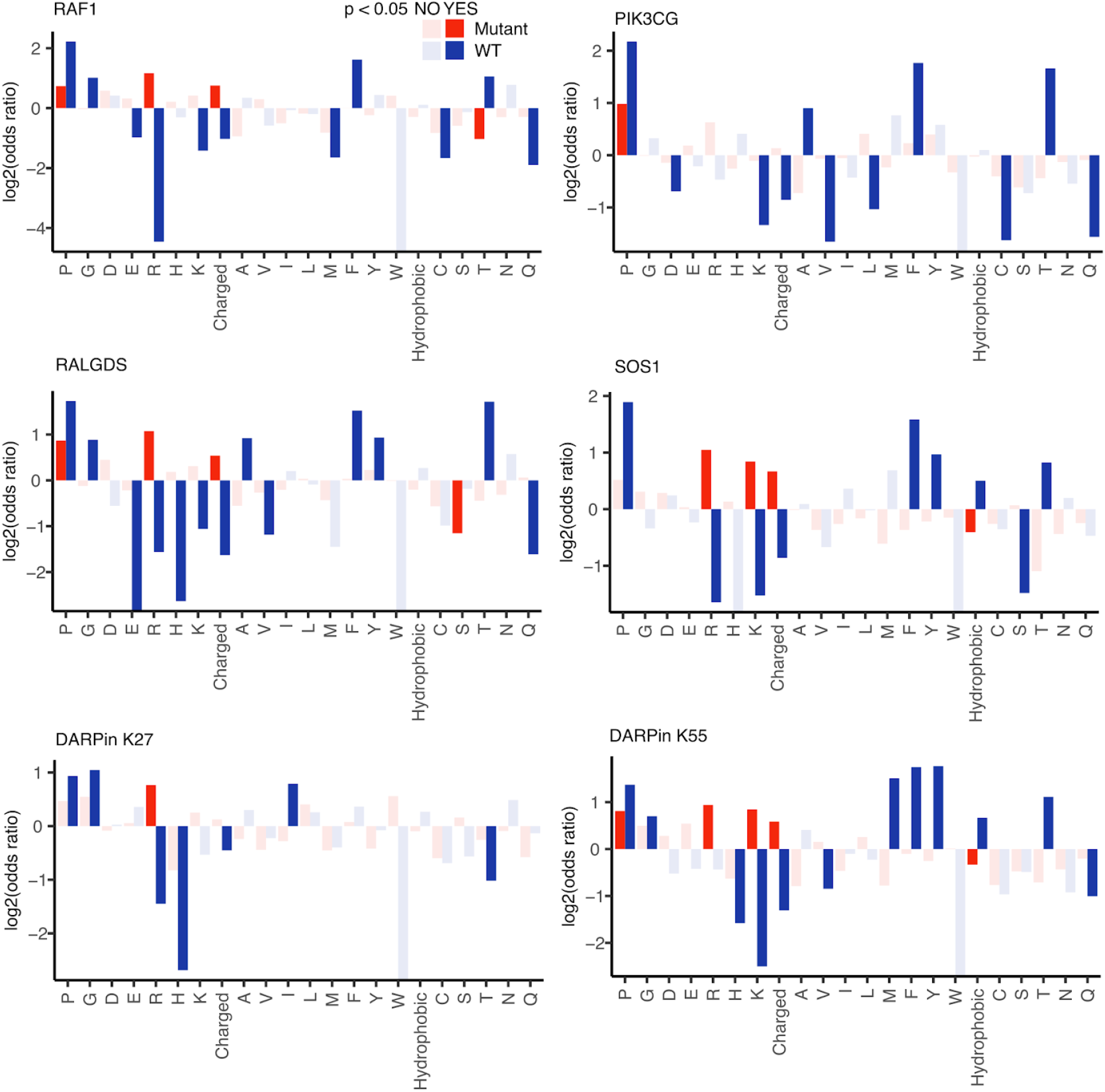
Enrichments of allosteric mutation types for each interaction. Enrichments are quantified for changes from each wild-type (WT) aa and for changes to each aa. Enrichments are also quantified for changes from and to amino acids with particular physicochemical properties: hydrophobic (A, V, I, L, M, F, Y, W) and charged (R, H, K, D, E). Results are shown for all residues outside the binding interface.

